# Deep learning enables the design of functional *de novo* antimicrobial proteins

**DOI:** 10.1101/2020.08.26.266940

**Authors:** Javier Caceres-Delpiano, Roberto Ibañez, Patricio Alegre, Cynthia Sanhueza, Romualdo Paz-Fiblas, Simon Correa, Pedro Retamal, Juan Cristóbal Jiménez, Leonardo Álvarez

## Abstract

Protein sequences are highly dimensional and present one of the main problems for the optimization and study of sequence-structure relations. The intrinsic degeneration of protein sequences is hard to follow, but the continued discovery of new protein structures has shown that there is convergence in terms of the possible folds that proteins can adopt, such that proteins with sequence identities lower than 30% may still fold into similar structures. Given that proteins share a set of conserved structural motifs, machine-learning algorithms can play an essential role in the study of sequence-structure relations. Deep-learning neural networks are becoming an important tool in the development of new techniques, such as protein modeling and design, and they continue to gain power as new algorithms are developed and as increasing amounts of data are released every day. Here, we trained a deep-learning model based on previous recurrent neural networks to design analog protein structures using representations learning based on the evolutionary and structural information of proteins. We test the capabilities of this model by creating *de novo* variants of an antifungal peptide, with sequence identities of 50% or lower relative to the wild-type (WT) peptide. We show by *in silico* approximations, such as molecular dynamics, that the new variants and the WT peptide can successfully bind to a chitin surface with comparable relative binding energies. These results are supported by *in vitro* assays, where the *de novo* designed peptides showed antifungal activity that equaled or exceeded the WT peptide.

## 1. Introduction

Proteins are one of the most interesting, widespread, and highly studied macromolecules due to their highly functional features. Proteins differ in their primary structure, which consists of a sequence of amino acids; this sequence strictly determines the spontaneous folding patterns and spatial arrangements that characterize their three-dimensional structures, which then define their diverse molecular functions^1,2^. Due to their diverse functional properties, proteins are of interest for a wide range of industrial applications. Rising interest from industry, combined with the increasing number of protein databases and structures and the development of more efficient computational algorithms, has led to the design of different pipelines for predicting the tertiary or quaternary structure and function of a protein from its primary structure^2^.

The vast diversity of protein structures is made possible by the effectively infinite number of possible combinations of twenty natural amino acids. This structural diversity has enabled the evolution of multiple functions responsible for most biological activities. Protein structures are usually determined by techniques such as NMR, X-ray crystallography, and cryo-electron microscopy, but *in silico* structure prediction methods have been shown to be an important alternative when experimental limitations exist. Protein structure prediction pipelines can follow a template-based or template-free approach (or a combination of both). In the template-based approach, protein structure decoys are built based on previously known protein structures^7-9^, whereas in the template-free approach, no structural templates are needed and new protein folds can be explored if desired^2^. One of the most widely-used approaches is fragment-based protein structure prediction. In this approach, decoys are built based on libraries of protein fragments with known structures (e.g. Rosetta^15-17^), with a search guided on angle torsions and secondary structures^2^.

Two general conventions exist for grouping proteins based on their structural similarity: i) homologous proteins inherit similarities from common ancestors, maintaining similar sequences and structures; and ii) analogous proteins have similar structures, given the limited local energy minima of their three-dimensional arrangements, but they do not necessarily maintain similar sequences with an evolutionary connection^3,4^. There are many examples of homologous proteins (i.e., proteins that share a common ancestor) as well as many tools for finding their evolutionary relationships^5,6^ and predicting their structure^7-9^. A classic example of protein homology is the TIM barrel fold, which is present in roughly 10% of all enzymes^10^. Moreover, homologous protein structures are widely classified in different databases such as CATH^12^, SCOP^13^, and Pfam^14^, while only one database exists for analogous motifs^3^. In general, fewer studies attempt to recognize structural analogs. Some examples of analogous structures are the hybrid motif βαβββ from the oligopeptide-binding protein OPPA in *Salmonella typhimurium* (PDB: 1B05), which is analogous to the core motif βαβββ from the antibiotic resistance protein FosA in *Pseudomonas aeruginosa* (PDB: 1NKI)^3^, and the artificial nucleotide-binding protein (ANBP, PDB: 1UW1), which is analogous to the treble clef zinc-binding motif^4,11^.

There are clear limitations to the available approaches for searching for structural analogs, and it has previously been suggested that more accurate statistical estimates are needed in order to identify similarities that are due to analogy rather than homology. It is therefore important to develop new alternatives that improve the conventional homology-based structure prediction approaches by also detecting analogous protein structures. One such alternative was presented by Zhang et al., who developed a pipeline for using docking-based domain assembly simulations to assemble multi-domain protein structures, with interdomain orientations determined based on the distance profiles from analogous protein templates^56^.

Recently, machine learning algorithms have received major recognition as an approach to predicting important sequence-structure relationships. Deep-learning (DL) strategies are neural networks with internal processing layers that can be trained to recognize patterns in large and complex data. DL strategies have been used for various protein applications, including the prediction of protein secondary structure and subcellular localization^19,20^; the prediction of protein contact maps, homology and stability^20^; protein design, such as the prediction of protein sequences based on protein structures^21^ and the design of metalloproteins^22^; and the prediction of protein folding^23,24^, among several other applications^25-30^. It is therefore of great interest to develop new tools that can accurately predict new protein sequence-structure relationships. Such tools will open doors for the automated search, prediction, and design of analog or low-homology proteins. These tools could additionally be used for other applications, such as the design of new protein structures and functions and the labeling of the dark proteome, in order to enrich scientific knowledge in these areas.

Recurrent neural network (RNN) models have already been used for protein domain prediction^41^. The information is provided to the model implicitly through large databases of unlabeled protein sequences (such as Pfam^42^) and of labeled structures (such as PDB and SCOPe^13^). These language models can extract biochemical and evolutionary information but lack structural information. Here we explore the capability of RNNs to capture structural information, including analogous structures to develop novel proteins. This alternative, based on Bepler’s model^41^, allows for the design of analog proteins with low sequence homology using a representation learning approach based on both evolutionary and structural information. To solve the lack of structural information, we stacked another model that allows the initial language model to learn structural information by predicting protein contact maps and secondary structures. By stacking these layers, we obtained a vector representation that allows us to find both structural and sequence similarities. Based on the similarity between these vector representations, we can search for and design proteins with desired structures and activities. Some of these structures and activities have not been explored by nature, given that we have not been able to find homologous proteins with more than 50% sequence identity. We demonstrate that this alternative model can capture both homology and analogy and could potentially improve template-based protein structure predictions, as well as other protein prediction tasks such as protein functionality.

In this work, we use this alternative protein design model to generate new *de novo* antifungal peptides. Antifungal peptides are receiving increasing interest due to their potential applications in diverse industries such as food manufacturing, agriculture, cosmetics, and therapeutics^36,37,38,39^, where they offer an attractive and useful replacement to current chemical alternatives.

## 2. Methods

### Language Model

Based on Bepler’s model^41^, we used a Bi-LSTM encoder as a language model (LM). The Bi-LSTM takes a protein sequence where each amino acid is a token (i.e., an instance of a sequence of characters in some particular document that are grouped together as a useful semantic unit for processing^40^) and represents the protein as a vector of the same length^41^. The LM architecture consisted of two Bi-LSTM layers with 1,024 hidden units in each layer, with a linear projection into the 20 amino acid prediction. We trained the model for six epochs with the ADAM optimizer, using a learning rate of 1*10^−3^ and a training batch size of 32 in a GPU V100. The LM was trained based on more than 21 million protein sequences obtained from Pfam^42^. We used the classical next-token prediction task with cross-entropy as a loss function.

### Structural Features Prediction

To predict protein contact maps and secondary structures as two different tasks, we stacked another two Bi-LSTM layers, each with 1,024 hidden dimensions and a projection, onto the initial LM. For the contact maps and secondary structure predictions, we used cross-entropy as a loss function. We trained the models for five epochs using the SCOP database^43^ filtered at 40% identity, which included more than 30,000 structures.

### Molecular dynamics simulations

Atomistic molecular dynamics (MD) simulations were run for the wild-type AC2 peptide (VGECVRGRCPSGMCCSQFGYCGKGPKYCGR, PDB: 1ZUV, with tryptophan 18 mutated back to phenylalanine), as well as for the two *de novo* design variants, DNv1-A C 2 (V Q D W C G N D C S A K E C C K R D G Y C G W G V D Y C G G) a n d D N v 2 - A C 2 (KRCGSQAGCPNGHCCSQYGFCGFGPEYCGR). Peptide models for DNv1-AC2 and DNv2-AC2 were based on the best matches from a homology search conducted using the HHpred bioinformatics toolkit^44,45^. The proteins that were used as templates were PDB 2KUS and 1MMC, which correspond to the antifungal peptides Sm-AMP-1.1a and AC2-WT, respectively. Mutations were introduced to these templates to achieve the desired sequences (shown above).

In the MD simulations, each of the three peptides was placed on top of, but not in direct contact with, a chitin surface that was formed by 14 polymers constructed with the *doglycan* software^46^ using the OPLS-AA force field^47^. Systems were solvated with water using the TIP3P model^48^ and then electro-neutralized with NaCl to a final concentration of 150 mM. For each peptide, two replicate simulations were run for 1 µs each. A leap-frog stochastic dynamics integrator was used to integrate Newton’s equations of motion with a time-step of 2 fs. Electrostatic interactions were calculated using the PME procedure^49^ with a real-space cut-off of 1.2 nm and a Fourier grid spacing of 0.12 nm. Van der Waals interactions were modeled using the classical Lennard-Jones potential with a cut-off of 1.2 nm. The LINCS^50^ algorithm was applied to constrain all H-bond lengths. Simulations were run at 1 atm with the Parrinello-Rahman barostat^51^ and at 298.15 K with the Berendsen thermostat^52^. Root mean square deviations in structure (RMSD) analyses were performed using GROMACS, with comparisons made against the crystal structure (or initial model) using Cα carbons only.

### MM/PBSA analysis for binding free energies

To calculate the peptide-chitin interaction energies, we used the MM/PBSA method as implemented using the default parameters in the software GMXPBSA 2.1^53^. Briefly, the interaction energy is represented as the sum of the molecular mechanics (MM) energy term and the Poisson-Boltzmann and surface area solvation (PBSA) term. The MM part is calculated as:

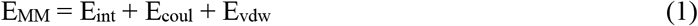

where E_int_ involves the bond, angle, and torsions and E_coul_ and E_vdw_ represent the electrostatic and Lennard-Jones energies, respectively. All these terms were extracted using GROMACS. For the PBSA part, the solvation term G_solv_ is composed of polar (G_polar_) and non-polar (G_nonpolar_) energy terms and is calculated as:

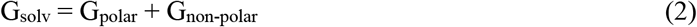

These terms were calculated using the Adaptive Poisson-Boltzmann Solver (APBS) software_54_. G_polar_ corresponds to the energy required to transfer the solute from a low dielectric continuum medium (ε=1) to a continuum medium with the dielectric constant of water (ε=80). In this case, we used the non-linearized Poisson-Boltzmann equation to calculate G_polar_. The G_nonpolar_ term is calculated as:

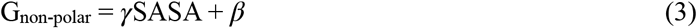

where = 0.0227 kJ·mol^-1^·Å^-2^ and *β*=0.0 kJ·mol^-1^. The dielectric boundary was defined using a probe of radius 1.4 Å. This protocol was performed for different “stable states “(see results section), where 100 frames spanning those regions were used for analysis.

### Microorganisms and growth media

*Escherichia coli* DH5α was used as the host for all DNA manipulations and vector storage and was grown in Luria-Bertani medium (LB: 10% tryptone, 5% yeast extract, 5% NaCl) supplemented with 100 µg/mL ampicillin (Amp) when needed. *E. coli* SHuffle® cells (New England Biolabs, USA) carrying our plasmids of interest were used for recombinant protein expression and were grown using Terrific Broth (TB: 1.2% yeast extract, 2.4% tryptone, 0.5% glycerol, 0.23% KH_2_PO_4_, 1.25% K_2_HPO_4_) supplemented with 2% glucose and 100 µg/mL Amp. The fungal species *Aspergillus niger, Fusarium oxysporum*, and *Trichoderma reesei* were routinely grown on Potato Dextrose Agar (PDA: 0.4% potato peptone, 2% dextrose, 1.5% agar). To obtain spores from the fungal species, PDA plates were seeded with fungal hyphae and grown at 30°C for one week. Spores were collected from the agar surface with sterile swabs, resuspended in sterile water, quantified by microscopy using a Neubauer chamber, and then stored at 4°C until use.

### DNA manipulation and cloning

Synthetic DNA fragments encoding the peptides AC2-WT, DNv1-AC2, and DNv2-AC2 were obtained from Integrated DNA Technologies (IDT, USA) and then amplified by PCR using primers that annealed at flanking attL sites. PCR products were cloned into the pETG41A plasmid by Gateway™ cloning using the LR Clonase II enzyme mix (Thermo Fisher Scientific, USA) following the manufacturer’s instructions, generating the plasmids pETG41A-AC2, pETG41A-DNv1, and pETG41A-DNv2. These vectors encoded the peptides of interest and were fused to maltose-binding protein (MBP) for increased solubility and a His6X tag for affinity purification. Plasmids were electrotransformed into *E. coli* DH5α. Successful transformants were selected on LB Amp plates; their plasmids were extracted by mini-prep and their constructs were confirmed by PCR and restriction assays. Verified plasmids were then electro-transformed into *E. coli* SHuffle® cells.

### Recombinant Protein Expression and Purification

*E. coli* SHuffle® cells carrying pETG41A-AC2, pETG41A-DNv1, and pETG41A-DNv2 were grown overnight in TB at 37°C with agitation. The following morning, 1 L flasks with 500 mL TB were inoculated with a 1:40 dilution of overnight cultures and grown at 37°C until an OD_600_ of ∼0.4 to 0.6 was reached. IPTG was then added to a final concentration of 0.1 mM, and cells were grown for 16 h at 20°C. Cells were collected by centrifugation, resuspended in lysis buffer (100 mM Tris-HCl, 300 mM NaCl, 5 mM Imidazole, 1 mM PMSF), and disrupted by sonication. Cell debris was removed by centrifugation for 40 min at 8500 rpm and 4°C. The soluble protein fraction was then purified with pre-equilibrated Ni-NTA resin (Qiagen), recovered in elution buffer (100 mM Tris-HCl pH 7.5, 300 mM NaCl, 350mM Imidazole and 10% glycerol), and dialyzed against protein storage buffer (100 mM TrisHCl pH 7.5, 300 mM NaCl, and 10% glycerol). Protein concentrations were determined using a 96-well based Bradford Assay (Bio-Rad, USA) in a BioTek Multi Plate Reader at A595 nm. Protein purity was checked by SDS-PAGE using SDS-12% polyacrylamide gel and Coomassie blue stain.

### Antifungal Activity Assays

To determine the minimum inhibitory concentration (MIC) of peptides in broth dilution assays, fungal spores were inoculated into Yeast Peptone Dextrose medium (YPD, 1% yeast extract, 2% peptone, 2% dextrose) at a concentration of 20,000 spores/ml. The inoculated medium was aliquoted into several tubes, and then serial two-fold dilutions of peptides were prepared. Tubes were incubated at 30°C for three days and visually inspected for the appearance of hyphal growth. To determine the MIC and IC_50_ of peptides in agar dilution assays, serial dilutions of antifungal peptides were prepared using protein storage buffer (as described above) and PDA plates were prepared by mixing one volume of tempered agar with one volume of peptide solution. Protein storage buffer was used as a negative control, and 100 µg/mL Zeocin (InvivoGen, USA) was used as positive antifungal control. We added 20,000 spores to the center of the plates before incubating plates at 30°C for at least 4 days, or until the negative control plate mycelium had grown to half the plate diameter. The diameter of fungal mycelium for each plate was measured and then normalized to the mycelium length of the negative control plate. The resulting data were plotted to interpolate IC_50_ using an asymmetrical sigmoid curve fit.

## 3. Results

### Embedding model and similarity metric

Bi-LSTM Recurrent Neural Networks (RNNs) can learn rich representations for natural language, which enables baseline performance on common tasks^55^. This model architecture learns by examining a sequence of characters in order and trying to predict the next character based on the model’s dynamic internal knowledge of the sequences it has seen so far (its “hidden state “). During the training phase, the model gradually revises the way it constructs its hidden state in order to maximize the accuracy of its predictions, resulting in a progressively better statistical summary, or representation, of the protein sequence. To examine what the model learned, we interrogated the model from the amino acid to the proteome level and examined its internal states. We then fine-tuned the language model using additional tasks, including secondary structure and contact map prediction. In total, the model processed ∼20,000 protein structures in ∼1 day on 2 Nvidia 2080 Ti GPUs (Fig. 1).

**Figure 1.**
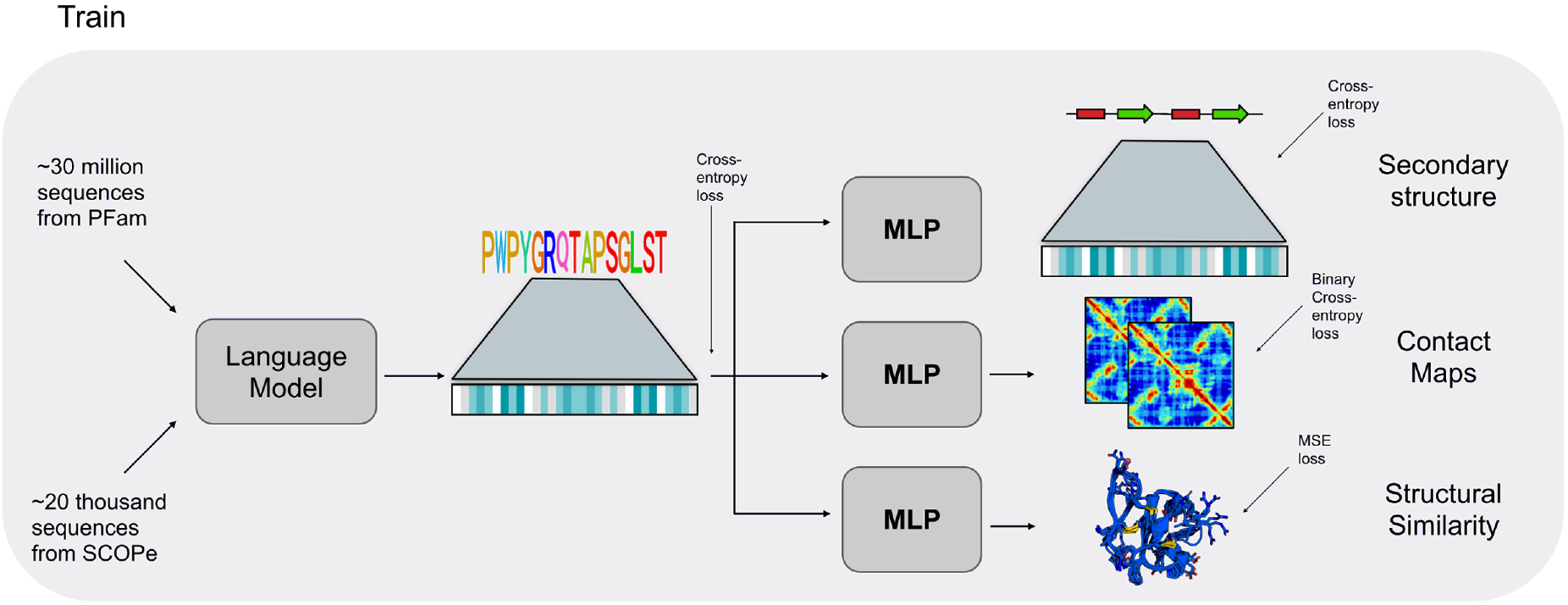
Training of the model. ∼21 million amino acid sequences from PFam and ∼20,000 sequences from the SCOPe database, encoding using amino acid character embeddings, were fed to the model. The model was trained to reconstruct a protein sequence while minimizing cross-entropy loss and then predict information about that sequence such as secondary structure, contact maps, and structural similarity.

The model used in this work could classify SCOP data (superfamily and fold classes) with 91.12% accuracy. We decided to study protein superfamily and fold classes because structural similarity between proteins remains challenging to infer solely from amino acid sequences. A previous study developed a method for encoding an amino acid sequence using structural information^41^. Using this method, any protein sequence can be transformed into a vector sequence encoding structural information, with one vector per amino acid position.

To assess the accuracy of our approach, we compared the structural similarity scores between proteins to the values obtained by comparing their sequences using this vector-encoding method. The first step in validating this alternative model was to validate the structural comparison model. A simple way to approach this problem would be to use metrics such as RMSD or dRMSD for structural comparison; however, these metrics need an equal number of elements to compare. The template modeling (TM) score measures the similarity of two protein structures, is more sensitive to the global fold similarity than to local structural variations, and is length-independent for random structure pairs^72^. Around ∼160,000 protein pairwise comparisons were therefore evaluated based on their TM scores. The results obtained from these pairwise comparisons corresponded with the structural similarity results obtained with the Bi-LSTM model, showing that an increase in the Bi-LSTM model energy criterion also meant a greater structural similarity according to the TM score (Fig. S2). These results show that protein pairs with a structural similarity score above 3.5 based on the SCOP hierarchy contain analogous structures, with TM scores ranging from 0.5 to 1.0.

### D*e novo* chitin-binding peptides from a language model

We used the similarity model to search for analogs of a chitin-binding protein and to design some *de novo* chitin-binding proteins. Starting from a sequence of poly-alanines, and based on what the language model had learned, we randomly mutated the peptide sequence until we obtained a structural similarity score greater than 3.5 (Fig. 2A). This threshold was based on the cut-off selected from Figure 2B. The mutation process took approximately 4 hours, during which the model explored approximately 20,000 possible sequences (Fig 3A). The mutation process was performed in three independent rounds, each with a different seed, in order to explore different model trajectories.

**Figure 2.**
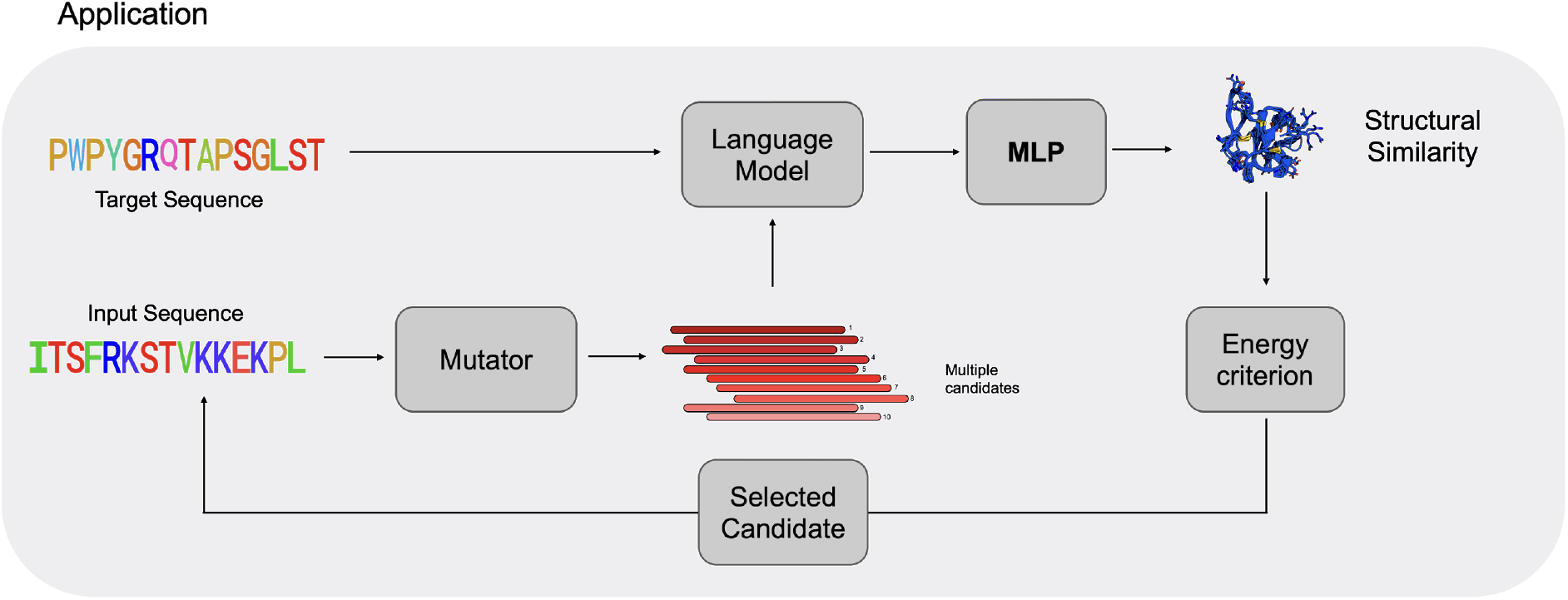
Application of the model. The model was used to generate *de novo* protein sequences by starting from a random or simple target protein sequence and iteratively mutating it to optimize a certain energy criterion.

**Figure 3.**
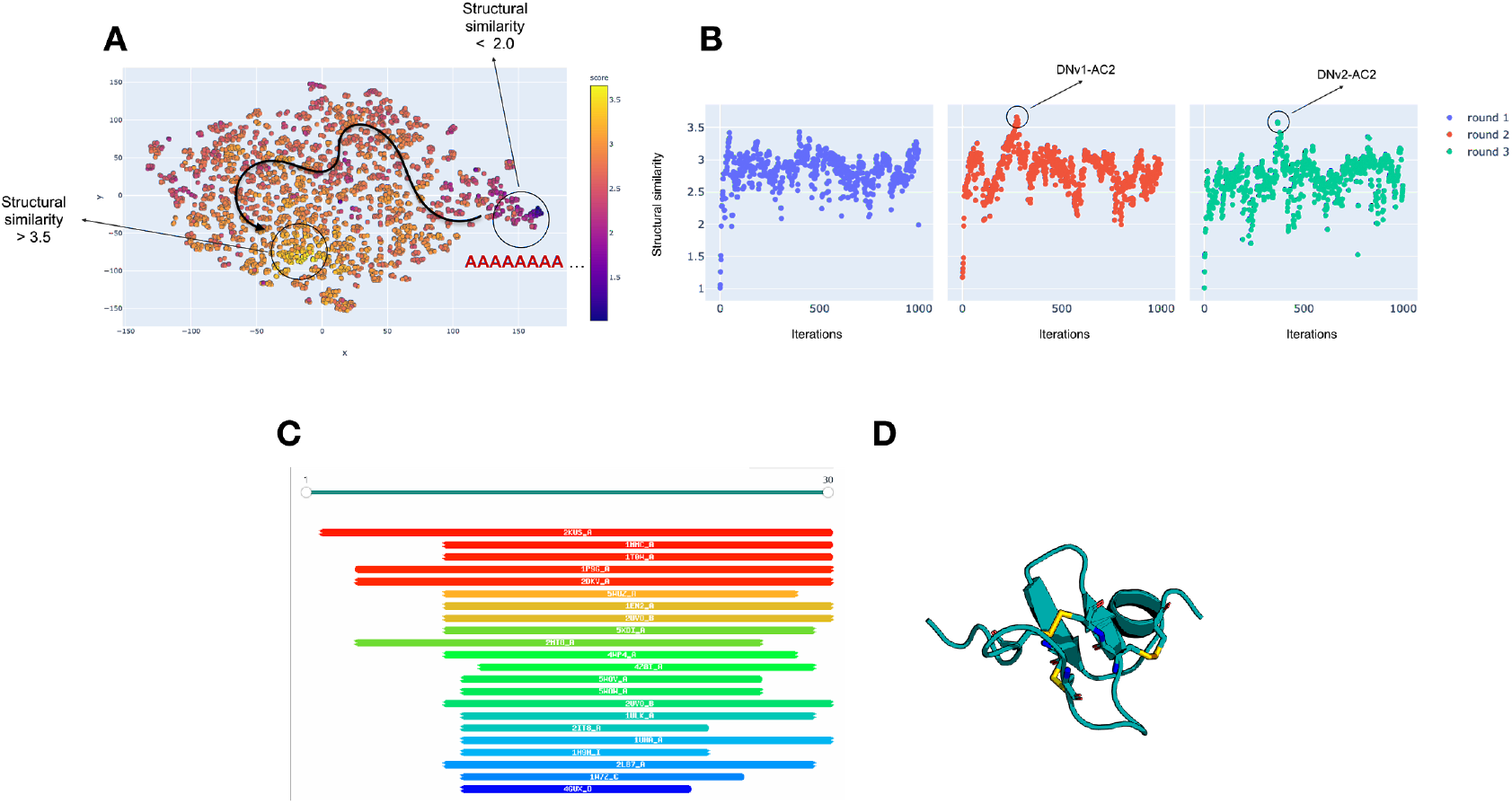
Design of the *de novo* protein variants. **(A)** Non-dimensional schematization of the *de novo* protein design pathway, which used the trained language model to implement a series of random mutations in an initial poly-alanine sequence (with an energy criterion below 2.0) in order to generate sequences with an energy criterion above 3.5. **(B)** Three independent trajectories were run for the *de novo* peptide design, and the two best candidates (DNv1-AC2 and DNv2-AC2) were selected for analysis. **(C)** HHPred search results for the peptide variants DNv1-AC2 and DNv2-AC2. **(D)** Three-dimensional model for DNv1-AC2 built using Modeller, based on the template protein 2KUS.

Interestingly, the two best *de novo* matches, DNv1-AC2 and DNv2-AC2 (Fig. 4B), shared just 40% and 54.8% sequence similarity, respectively, with the AC2-WT variant, but were still predicted to have chitin-binding activity. For both of the *de novo* peptides, the first homologous match obtained by HHpred^44,45^ was to PDB 2KUS, which corresponds to the antimicrobial peptide Sm-AMP-1.1a, an antifungal peptide with a chitin-binding domain (Fig. 3C and Fig. S1). A second match, with a similar score, was to PDB 1MMC, which corresponds to AC2-WT (Fig. 3C and Fig. S1). The sequence percentage identity for the *de novo* peptides DNv1-AC2 and DNv2-AC2 against the 2KUS match was 32.4% and 48.6%, respectively. All of the presented sequence identities do not pass the threshold for structurally reliable homologous alignments, which is determined as a function of the alignment length^57^ (which in this case is 30 residues). These peptides could therefore be considered analogous. The two HHpred matches were used for the construction of peptide models (Fig. 3D) and for the following structural analyses of the designed peptide variants.

**Figure 4.**
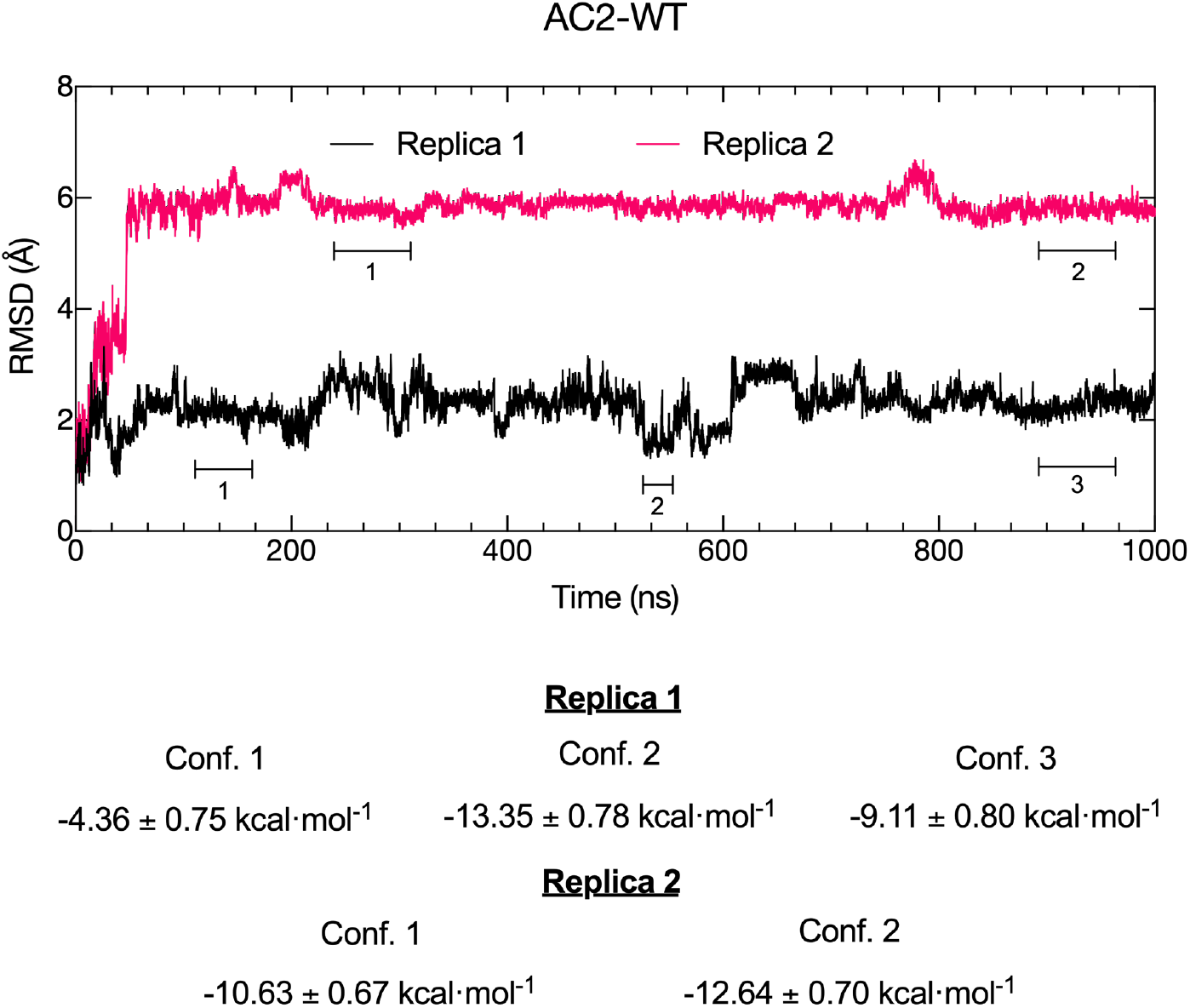
RMSD and relative binding energies for AC2-WT. RMSD time-series data for AC2-WT across two replicate simulations. Different configurations were obtained at different simulation times, and an MM/PBSA calculation was performed for each configuration. Relative binding free energies are shown at the bottom for each of the configurations depicted in the RMSD plot for each replica.

### Molecular dynamics and interactions with a chitin surface for AC2-WT and the AI-generated peptide variants DNv1-AC2 and DNv2-AC2

#### 1. AC2-WT2

To preliminarily explore the molecular interactions of the tested antifungal peptides (AC2-WT, DNv1-AC2, and DNv2-AC2), we performed unbiased MD simulations of a single free peptide near a chitin surface (Fig. S3). Two replicate simulations were run for 1 µs each for each peptide variant. We used MM/PBSA calculations to evaluate both the potential for spontaneous binding of the peptides to the chitin surface as well as the strength of the peptide-chitin interaction.

We first evaluated these simulations for the AC2-WT antimicrobial peptide. RMSD time-series data showed important deviations between the simulated AC2-WT structure and its crystal structure (Fig. 4). The highest RMSD values that replica 1 achieved were approximately ∼3.0 Å, whereas replica 2 achieved values near ∼6 Å, when the peptide was compared with its crystal structure (Fig. 4). Both simulations achieved a roughly stable RMSD after the first 50 ns. Interestingly, despite the high RMSD for replica 2, the predicted relative binding energy for replica 2 was either higher (−10.63 to -12.64 kcal·mol^-1^) than some of the selected structural configurations from replica 1 (−4.36 and -9.11 kcal·mol^-1^ for configurations 1 and 3 in replica 1, respectively) or within the same order of magnitude (−13.35 kcal·mol^-1^ for configuration 2 in replica 1) (Fig. 4).

The selected AC2-WT protein configurations with residues that were in contact with the chitin surface are depicted in Figure 5. In replica 1, the number of interactions increased over time, with more residues seen close to the chitin surface (Fig. 5). However, more peptide-chitin contacts did not translate to higher interaction energy, since the interaction energy changed from -13.35 to -9.11 kcal·mol^-1^ between configurations 2 and 3 (Fig. 4 and Fig. 5A). In replica 2, most of the peptide-chitin contacts stabilized earlier in the simulations, with no important visual differences (Fig. 5A). Despite the differences among replicas and configurations, there was an observable pattern in the regions in contact with the chitin surface (Fig. 5B), with the beginning, middle, and end of the protein structure involved in most of the contacts. The only difference between replicas was that the initial region of the AC2-WT peptide stayed in contact with the chitin surface in replica 2 but not in replica 1 (Fig. 5B). It is also important to note that AC2-WT interacted with the chitin surface through aromatic residues F18, Y20, and Y27. Aromatic residues have been previously classified as conserved key residue interactions for the hevein-like peptides mechanism^58-60^.

**Figure 5.**
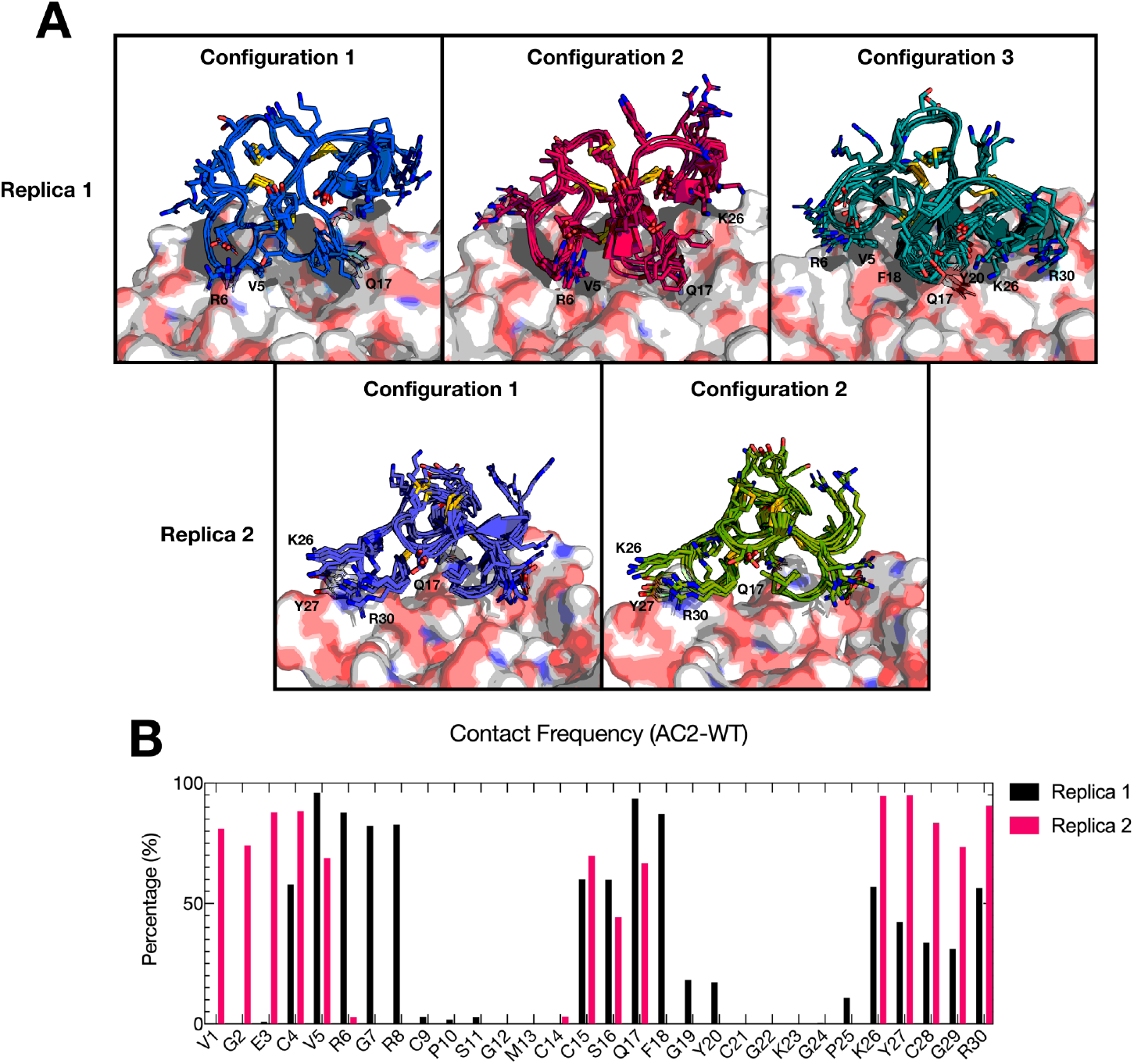
Contact analyses for AC2-WT. **(A)** Sample snapshots from the configurations used in the analyses of binding energies for AC2-WT (replicas 1 and 2). Residue numberings are shown for the residues that are visually close to the chitin surface. **(B)** Per-residue percentage of contact for AC2-WT residues across the entire simulation, for both replica 1 and 2, using a distance cut-off of 4.5 Å.

#### 2. DNv1-AC2

Similar analyses were performed for the AI-generated peptide variant DNv1-AC2. RMSD time-series data showed that the peptide conformation remained close to the initial conformation in both replicas, with average RMSD values of 2.42 and 3.68 Å for replicas 1 and 2, respectively (Fig. 6). These roughly stable conformations were achieved early, after approximately 30 ns of the simulation, which was similar to the AC2-WT simulations (Fig. 6). In terms of binding free energies, the two selected configurations from replica 1 achieved energies of the same magnitude as AC2-WT, with values of -14.47 kcal·mol^-1^ and -13.40 kcal·mol^-1^ (Fig. 6). The contact regions in replica 1 of the DNv1-AC2 variant are similar as well, with the main contacts located in the middle and end portions of the structure (Fig. 7A and 7B) and some contacts locating at the beginning, especially at residue G6, which was in contact with the chitin surface for 83% of the simulation (Fig. 7B).

**Figure 6.**
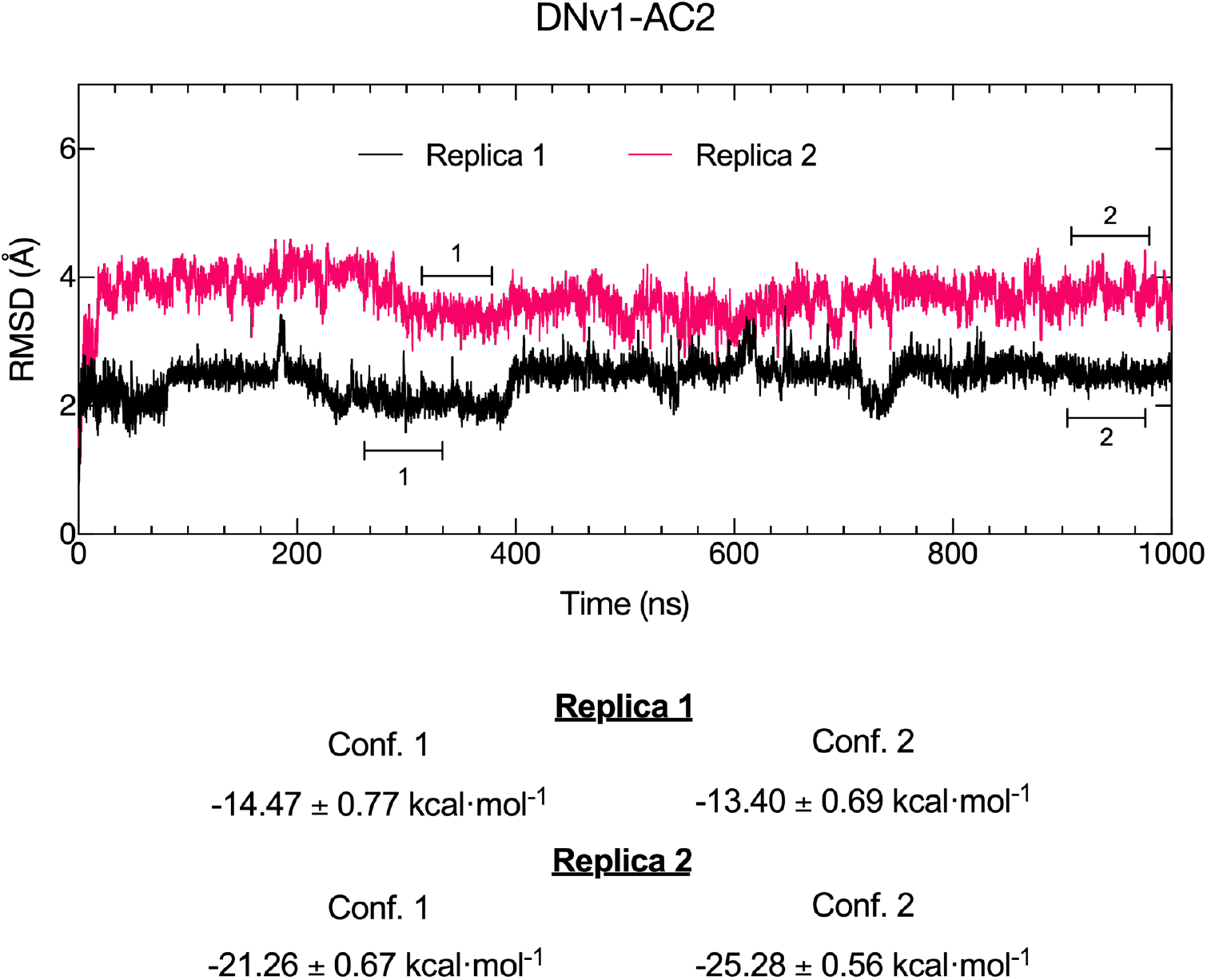
RMSD and relative binding energies for DNv1-AC2. RMSD time-series data for the DNv1-AC2 peptide variant across two replicate simulations. Different configurations were obtained at different simulation times, and an MM/PBSA calculation was performed for each configuration. Relative binding free energies are shown at the bottom for each of the configurations depicted in the RMSD plot for each replica.

**Figure 7.**
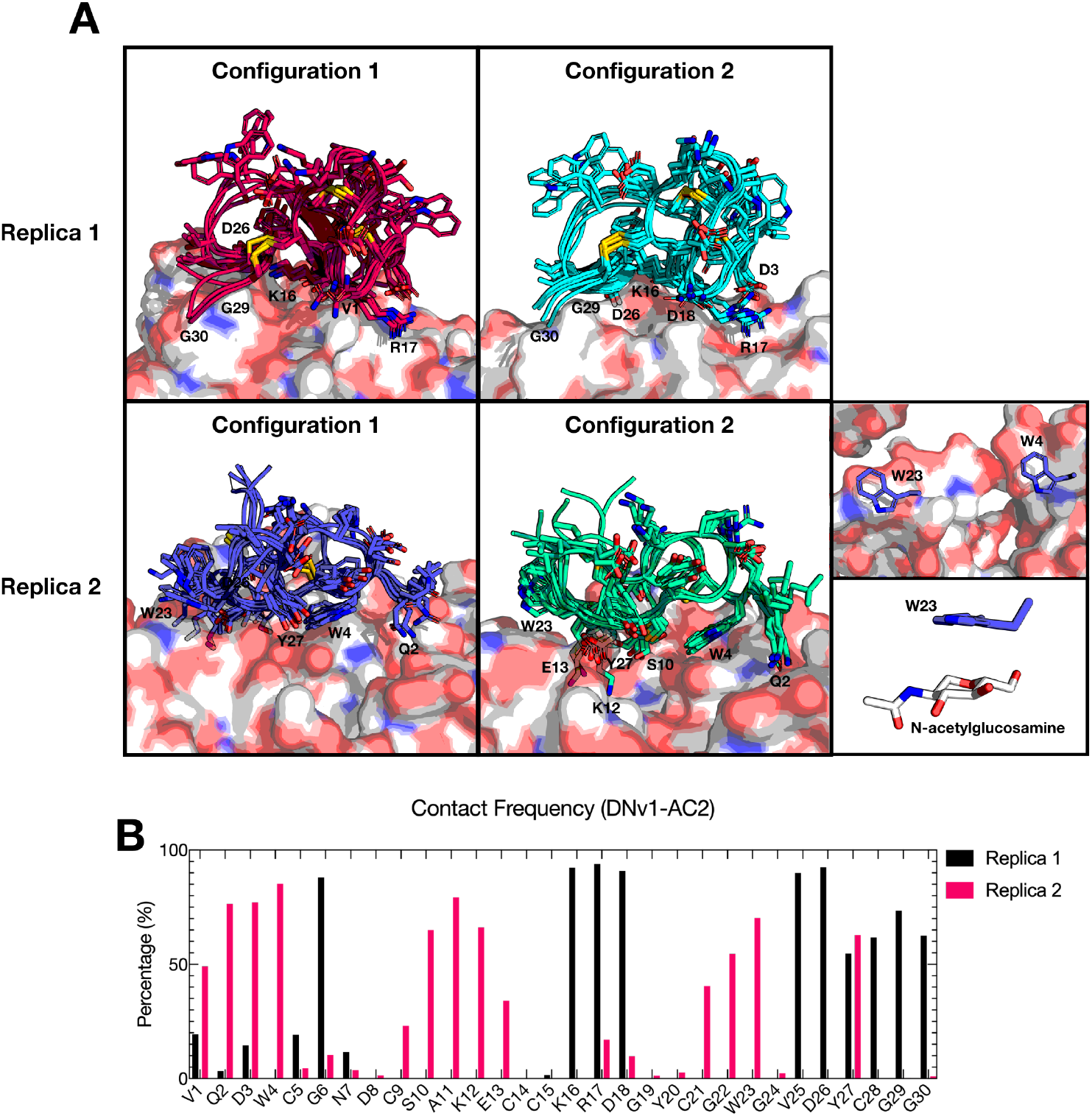
Contact analyses for DNv1-AC2. **(A)** Sample snapshots from the configurations used in the analyses of binding energies for the DNv1-AC2 variant (replica 1 and 2). Residue numberings are shown for the residues that are visually close to the chitin surface. **(B)** Per-residue percentage of contact for DNv1-AC2 residues across the entire simulation, for both replica 1 and 2, using a distance cut-off of 4.5 Å.

Interestingly, replica 2 behaved differently from replica 1. The relative binding energies were higher for both selected configurations from replica 2, with values of -21.26 kcal·mol^-1^ and -25.28 kcal·mol^-1^ (Fig. 6). The contact area was different from replica 1 as well, with most contacts at the beginning (from residue 1 to 7) and middle (from residue 9 to 13) of the structure (Fig. 7A and 7B). New contacts were also established in the region between residues 21 and 23 (Fig. 7A and 7B). One possible explanation for the higher energies of this configuration is the interactions of residue W4 and W23, as shown in Figure 7A (bottom right panels). It has previously been shown that these types of interactions, where tryptophan residues are flatly aligned in a CH-π orientation, occur between chitinases and chitin. These interactions are also frequently observed between proteins and sugars^61^. Previous data supports this generalization and suggests that larger aromatic groups have higher association constants and binding enthalpies^62^. In addition, residue E13 can form hydrogen bonds with the amide group of N-acetylglucosamine, which could enhance its ability to bind to the chitin surface.

#### 3. DNv2-AC2

RMSD time-series data for the DNv2-AC2 variant showed similar results to the DNv1-AC2 variant, with values of 3.94 Å and 3.24 Å for replica 1 and 2, respectively (Fig. 8). Two conformations were selected from each replica, spanning the simulation times shown in Figure 8, and MM/PBSA calculations were performed for each conformation. As shown in Figure 8, the relative binding energies were dependent on the peptide configuration. For replica 1, the first selected conformation only exhibited an average energy of -7.47 kcal·mol^-1^ (Fig 8), whereas the binding energy increased to -33.49 kcal·mol^-1^ by the second configuration. The main difference between these two configurations is the presence of aromatic residues, such as residues Y18, F20 and Y27, in direct contact with the chitin surface in configuration 2. These residues are similar to the AC2-WT peptide (Fig. 9A), and, as previously mentioned, seem to be important for the peptide’s activity^58-60^.

**Figure 8.**
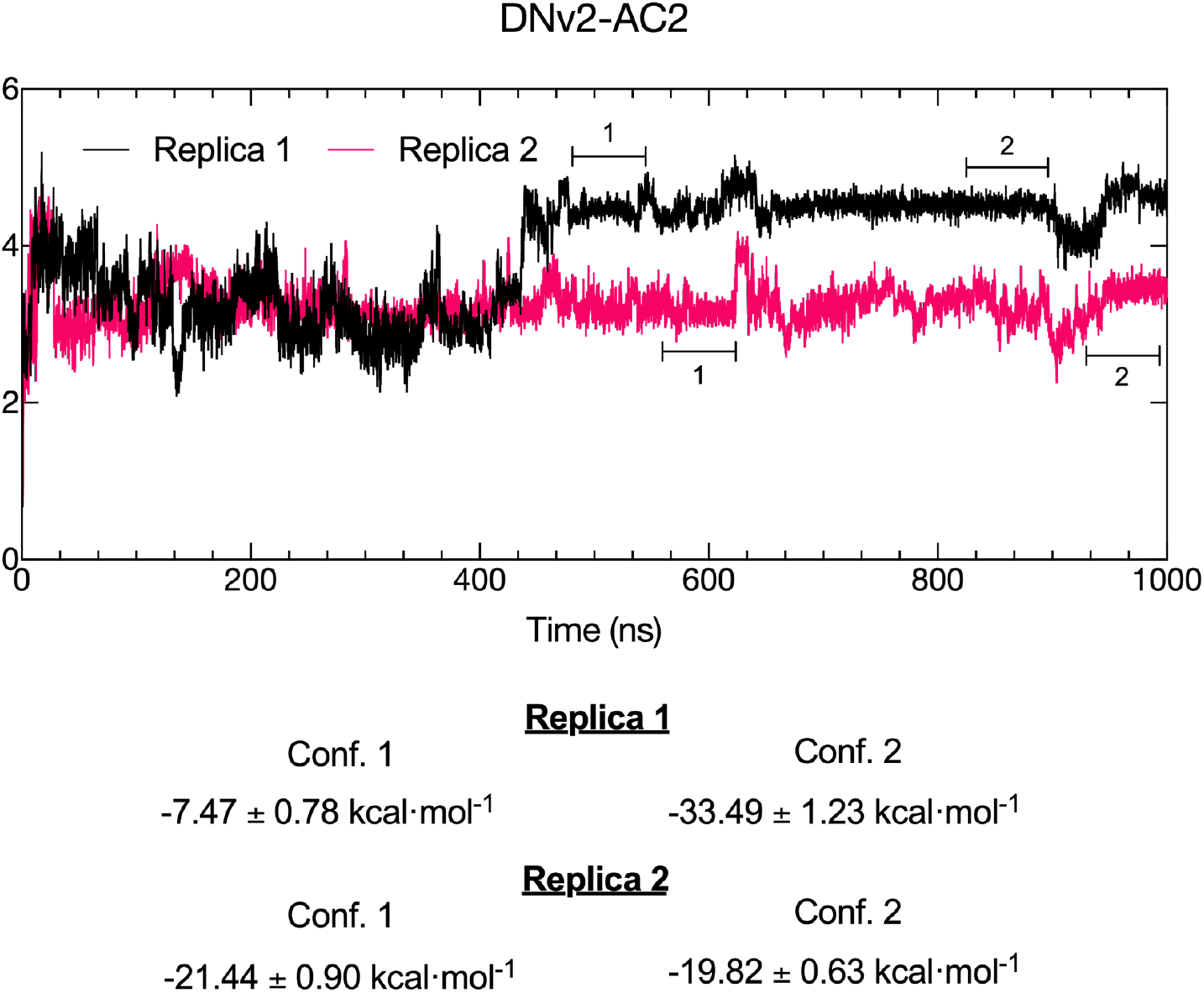
RMSD and relative binding energies for DNv2-AC2. RMSD time-series data for the DNv2-AC2 peptide variant across two replicate simulations. Different configurations were obtained at different simulation times, and an MM/PBSA calculation was performed for each configuration. Relative binding free energies are shown at the bottom for each of the configurations depicted in the RMSD plot for each replica.

**Figure 9.**
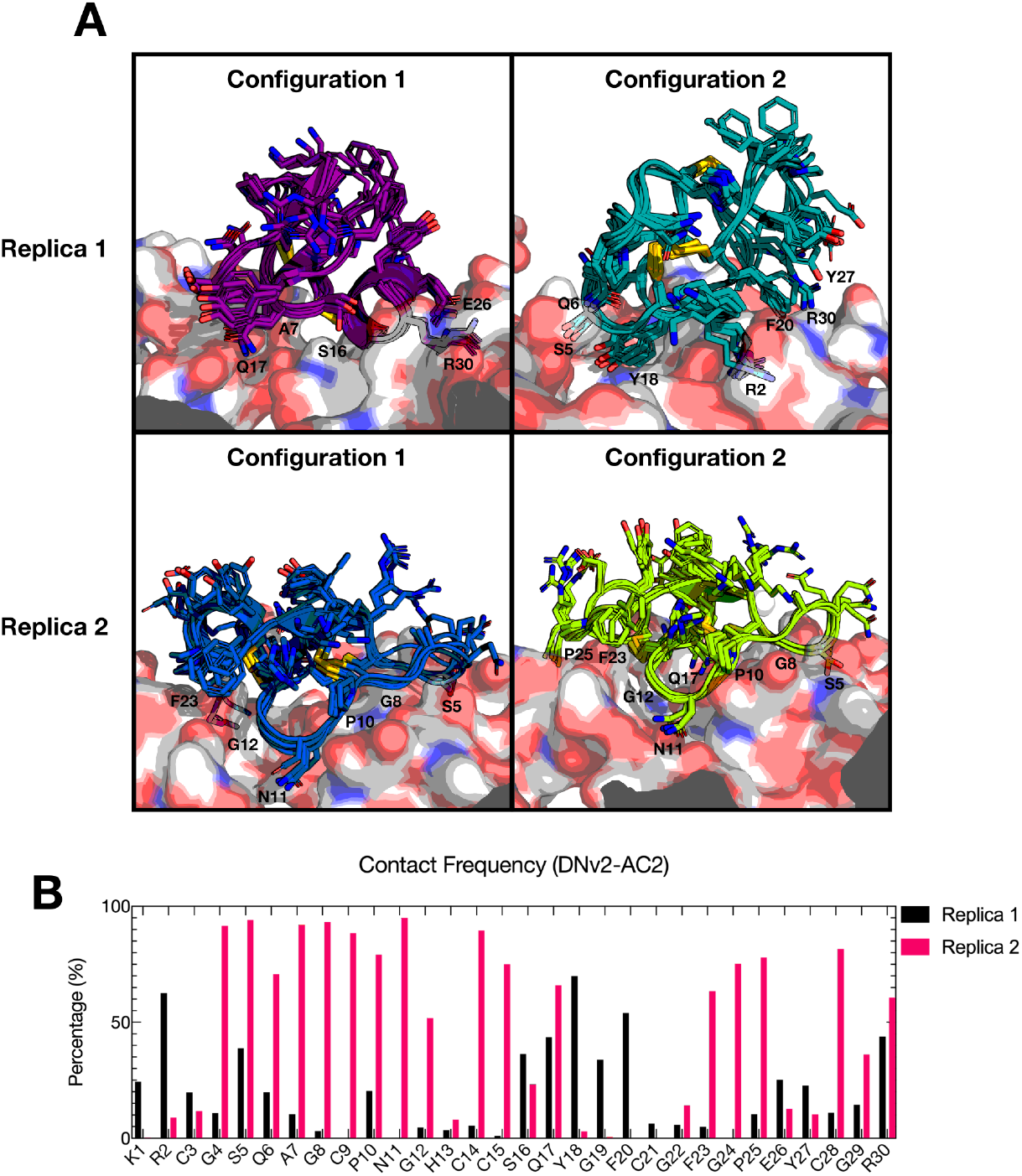
Contact analyses for DNv2-AC2. **(A)** Sample snapshots from the configurations used in the analyses of binding energies for the DNv2-AC2 variant (replicas 1 and 2). Residue numberings are shown for the residues that are visually close to the chitin surface. **(B)** Per-residue percentage of contact for DNv2-AC2 across the entire simulation, for both replica 1 and 2, using a distance cut-off of 4.5 Å.

In replica 2, the peptide-chitin interactions occurred mostly at the hydrophobic residues G4, A7, G8, G12, G24, and G29. Residue F23 was the only aromatic residue in direct contact with the chitin surface (Fig. 9A). These differences may explain the strength of the interaction compared to the second configuration from replica 1 (Fig. 8). However, this peptide-chitin interaction is still stronger than AC2-WT and is similar to the DNv1-AC2 variant, potentially due to residues Q6, Q17, and R30, which can help in the formation of salt bridges and hydrogen bond interactions (Fig. 9). It is important to note that arginine residues, which were present in all the tested variants and were also seen directly interacting with the chitin surface (Figs. 5, 7, and 9), allow the peptides to interact with anionic components. These interactions also help in the formation of salt bridges and hydrogen bonds. Like tryptophan, arginine can participate in cation–π interactions, which enhances the interactions between peptides and their targets^63^.

### *De novo* peptides exhibit *in vitro* inhibitory activity against different fungal species

We validated the DNv1-AC2 and DNv2-AC2 *de novo* peptide designs by testing their antifungal activity and potency relative to the AC2-WT peptide *in vitro*. DNA encoding the DNv1-AC2, DNv2-AC2, and AC2-WT peptides was cloned into expression vectors to allow recombinant production of the WT peptide and *de novo* designs in *E. coli*. All peptides were expressed as chimeric fusions to maltose-binding protein (MBP) in order to improve their solubility and to facilitate purification and downstream handling. All constructs showed a good level of protein production after vector expression was induced and were purified for downstream functional characterization (Fig. 10A).

**Figure 10.**
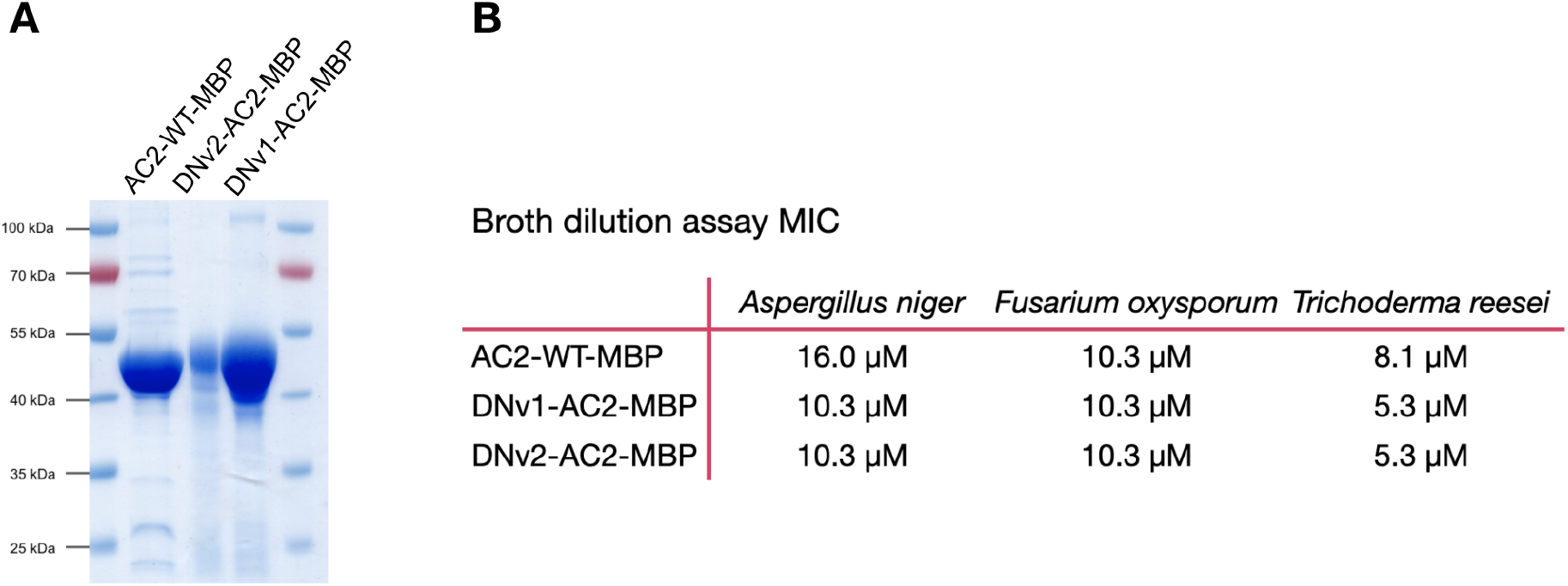
Purification and activity of WT and *de novo* AC2 peptides fused to MBP protein. **(A)** SDS-PAGE stained with Coomassie blue dye, showing purified peptide-MBP fusion proteins. **(B)** Evaluation of the antifungal activity of peptide-MBP fusion proteins. Broth dilution assays were performed to determine the minimum inhibitory concentration (MIC) of each peptide against three fungal species. Each peptide-MBP dilution in broth media was tested in duplicate, and growth was assessed visually.

We first used a broth dilution assay to test the activity of the peptide-MBP fusions against three filamentous fungi: *Aspergillus niger, Fusarium oxysporum*, and *Trichoderma reesei* (Fig. 10B). AC2-WT-MBP had antifungal activity against all three tested fungi, with MICs ranging from 8 to 16 µM. These results corroborate the phenotype described for the native Ac-AMP2 protein purified from *Amaranthus caudatus* grains^64^. DNv1-AC2-MBP and DNv2-AC2-MBP both exhibited similar inhibitory activity to AC2-WT when tested against *F. oxysporum*. Surprisingly, both *de novo* peptides showed slightly increased activity against *A. niger* and *T. reesei* compared to AC2-WT-MBP.

To further investigate the performance of the *de novo* peptide-MBP fusions, we used agar dilution assays to test for the inhibition of *A. niger*. The differences in the antifungal potency of both peptides was readily apparent in these assays. As seen in Fig. 11A, DNv1-AC2-MBP slightly decreased mycelium growth at lower concentrations and greatly inhibited growth at concentrations greater than or equal to 7.5 µM. In comparison, DNv2-AC2-MBP negatively impacted mycelium growth even at low micromolar concentrations. The aggregated results of the different experiments and replicates are summarized in Figure 11B, with IC_50_ values of 5.2 µM for DNv1-AC2-MBP and 2.5 µM for DNv2-AC2-MBP, indicating a higher potency of the latter peptide variant.

**Figure 11.**
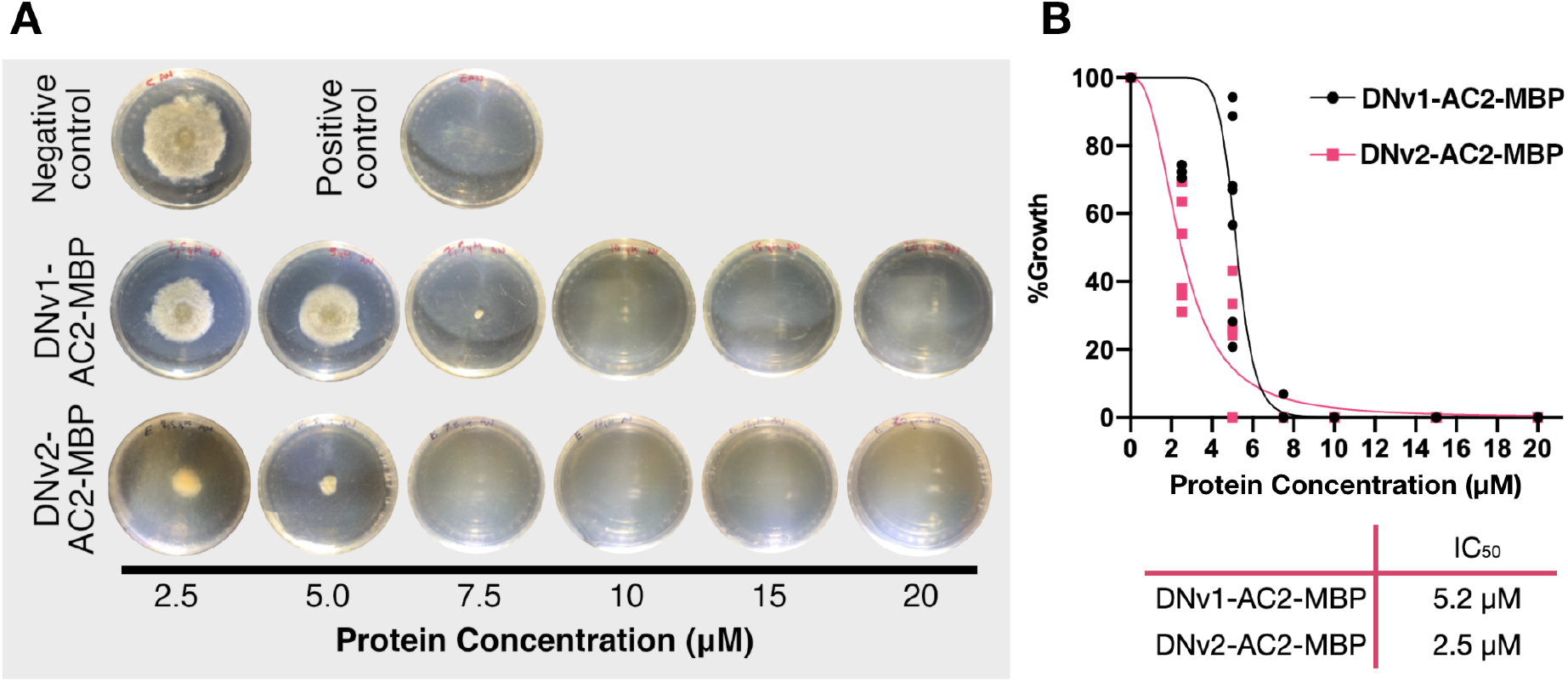
Agar dilution assay comparing the antifungal activity of DNv1-AC2-MBP and DNv2-AC2-MBP against *A. niger*. (A) Representative images of agar dilution assays performed with serial dilutions of *de novo* peptide-MBP fusions added to agar before solidification. Assays tested for the inhibition of mycelial growth from *A. niger* spores. Protein storage buffer was used as a negative control for growth inhibition, and Zeocin was used as positive control. (B) IC_50_ values for DNv1-AC2-MBP and DNv2-AC2-MBP. Aggregated results from three independent experiments were used to interpolate IC_50_ from the fitted curve.

Altogether, these results show that our *de novo* designed peptides replicated the antifungal activity of the AC2-WT protein, and, in the case of DNv2-AC2, did so with higher potency against the target fungi. These *in vitro* results confirm the results obtained by molecular dynamics simulations and validate our alternative model, based on previous RNNs, for *de novo* protein design.

## 4. Discussion and Conclusions

In this work, we developed and trained a deep-learning model, based on previous RNNs, and showed how it can be used to guide and improve protein design. Specifically, our alternative approach can design analog proteins based on a representation learning architecture that contains evolutionary and structural information from millions of protein sequences. Given that the study of sequence-structure relationships in proteins is a highly dimensional task, this type of approach can play an important role in this area and complement similar models^30,41,58,59^. We have shown that this alternative approach promises high-quality predictions, with an accuracy of 91.12% when classifying structural information from the SCOP database^41^. These results are similar to other models available in the literature^58,59^. We have additionally shown that it is possible to generate *de novo* protein sequences with a particular function; in this work, we demonstrated this by generating peptides with antifungal activity. Interestingly, the model can generalize, as shown by its ability to successfully generate functional proteins starting from a simple poly-alanine sequence. To put the capabilities of the model in context, we extended our study of the predicted *de novo* antifungal peptides by running molecular dynamics simulations, evaluating the interactions involved in the process of chitin recognition, and observing patterns such as the importance of aromatic residues to the strength the binding interactions.

To date, numerous methods have been described for generating novel antifungal peptide designs. Directed and rational approaches have focused on comparing the sequences and structures of antifungal peptides to identify the key elements that impact antifungal potency, such as peptide cationicity and hydrophobicity, peptide tertiary structure and the distribution of the residues within that structure, and peptide length and amphipathicity^36,65,66^. On the other hand, machine learning methods have become an attractive alternative for predicting peptide sequences with antifungal activity; different publications have explored this approach by using and combining a variety of approaches like support vector machines^67^, hidden Markov models (HMM)^68,69^ and character embedding^70^. In this work, our alternative deep-learning model, based on previous RNNs^41^, generated *de novo* peptide sequences with potent *in vitro* antifungal activity that was comparable to wild-type antifungal variants such as AC2-WT. In the case of DNv2-AC2, the *de novo* peptide even showed improved potency compared to its native counterpart, highlighting the power of this approach to generate novel and useful peptide variants.

The results presented in this work open a huge door for the development of new alternative proteins and peptides, and the model presented here has potential applications in industries such as food manufacturing, agriculture, cosmetics, and therapeutics, as well as in the design of new proteins with specific activities. We also believe that, by providing an example of the artificial generation of functional protein analogs, this manuscript broaches an important topic and encourages discussions of simplicity and limits in nature, especially in terms of the possible structural arrangements a protein can adopt. Homology is the classical definition for similarity at the sequence, structure and function levels, but no clear definition exists for considerable structural similarities despite low sequence identities. The term “remote homology “has been used to describe similar structures with a sequence identity of 25% or lower; this type of homology is usually inferred from common features, such as functional residues, or from unusual structural features^71^. On the other hand, the term “analogy “usually refers to two or more proteins with no common origin that converge to similar structural features^4^. Both cases are difficult to experimentally validate and their definitions have changed throughout the years. Finally, our approach has certain advantages over the state-of-art models, such as its low complexity (i.e., fewer parameters) and its inclusion of structural information. Limitations of these types of models are related to the size variability of the generated embeddings, which makes it difficult for comparison and interpretability.

## Supplementary information

**Fig. S1.**
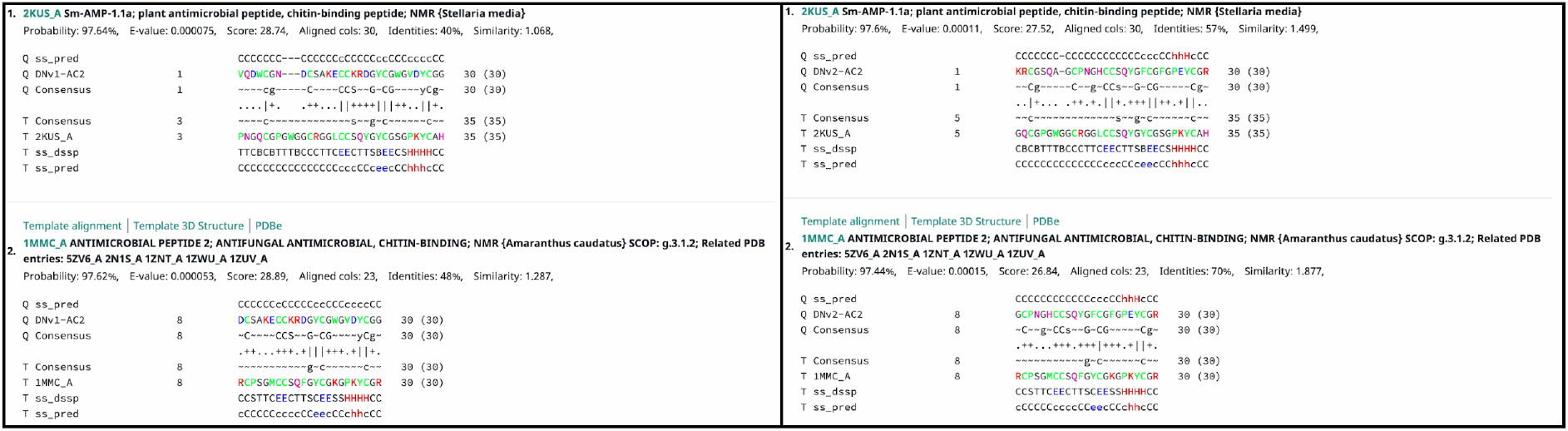
HHpred search results for *de novo* variants. First and second matches for the *de novo* variants DNv1-AC2 (left) and DNv2-AC2 (right), found with the HHpred bioinformatics toolkit. Matches correspond to the antimicrobial peptides Sm-AMP-1.1a (PDB: 2KUS) and AC2-WT (PDB: 1MMC).

**Fig. S2.**
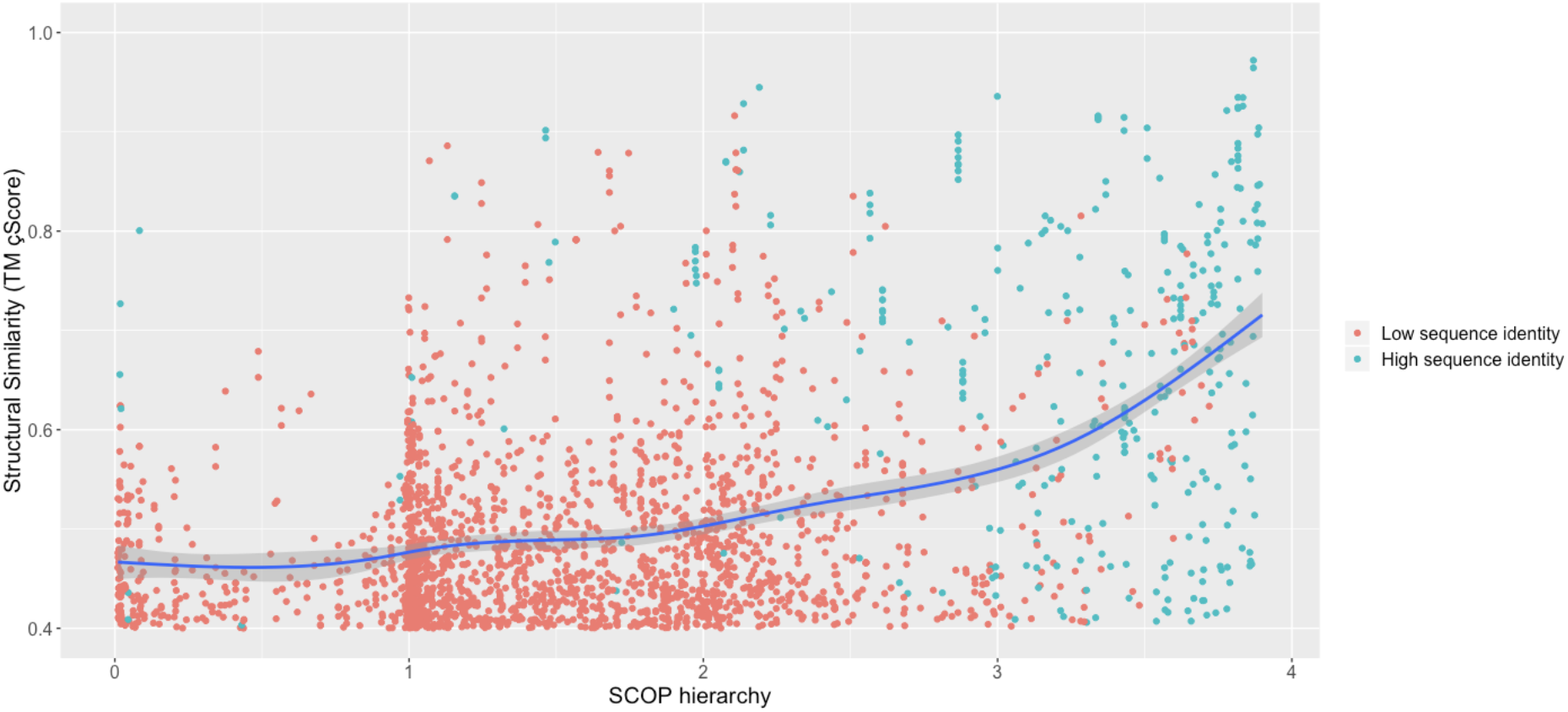
Correlation between TM score and predicted SCOP hierarchy. All 160,000 pairwise comparisons show a positive correlation when compared with the structural similarity score (TM score). Plot show only pairwise comparisons with a TM score above 0.4.

**Fig. S3.**
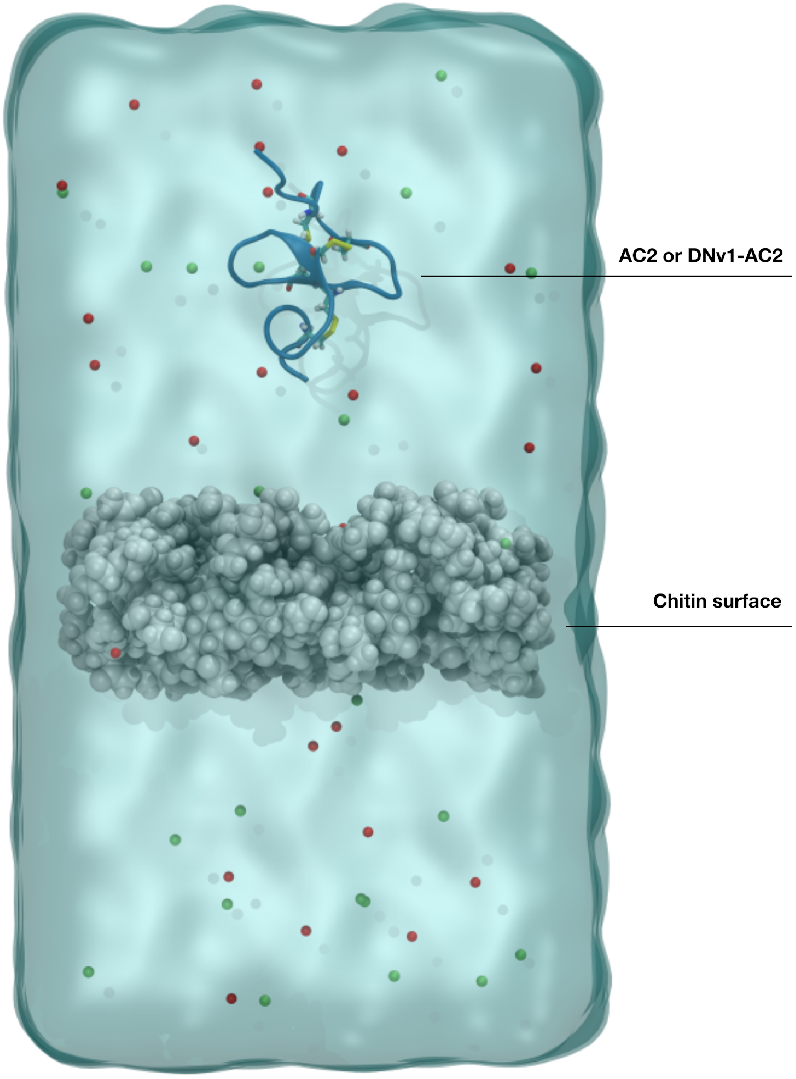
Schematic representation of the simulated systems. The system setup was similar for both antifungal peptide variants, AC2-WT or DNv1-AC2. A chitin surface was built using 14 polymers formed by 7 N-acetylglucosamide monomers. The system was solvated in water, electro-neutralised in NaCl (red and green spheres), and left at a concentration of 150 mM.

